# Independent respiratory and locomotor rhythms in running mice

**DOI:** 10.1101/2020.08.09.242768

**Authors:** Coralie Hérent, Séverine Diem, Gilles Fortin, Julien Bouvier

**Affiliations:** Université Paris-Saclay, CNRS, Institut des Neurosciences Paris-Saclay 91190, Gif-sur-Yvette, France; Institut de Biologie de l’École Normale Supérieure (IBENS), École Normale Supérieure, CNRS, INSERM, PSL Research University, 75005 Paris, France

**Author notes:** Corresponding author information: Université Paris-Saclay, CNRS, Institut des Neurosciences Paris-Saclay 91190, Gif-sur-Yvette, France.

## Abstract

Examining whether and how the rhythms of limb and breathing movements interact is highly informative about the mechanistic origin of hyperpnoea to exercise. However, studies have failed to reveal regularities. In particular, whether breathing frequency is inherently proportional to limb velocity and/or imposed by a synchronization of breaths to strides is still unclear. Here, we examined the specifications of respiratory changes during running in mice, the premier model for investigating, in a standardized manner, complex integrative tasks including adaptive breathing. We show that respiratory rate increases during running to a fixed and stable value, irrespective of trotting velocities and of inclination. Yet, respiratory rate was further enhanced during gallop. We also demonstrate the absence of temporal coordination of breaths to strides at any speed, intensity or gait. Our work thus highlights a hardwired mechanism that sets respiratory frequency independently of limb movements but in relation with the engaged locomotor program.

## INTRODUCTION

The versatile adaptability of breathing to changes in the environment or behavioral state is vital. Probably the most striking example is the augmentation of ventilation at the transition from rest to running exercise to match the augmented energetic demand (Bramble and Carrier, 1983; Mateika and Duffin, 1995; Gariepy et al., 2010). The hyperpnoea during running is principally supported by an increased respiratory rate which underscores an upregulation of the respiratory rhythm generator in the brainstem (Del Negro et al., 2018). While examining the dynamic interactions between respiratory and locomotor movements should inform on the origin and nature of the activatory signal, studies have failed to reveal regularities. Despite this, a common postulate is that respiratory frequency is entrained by that of locomotor movements, through inertial oscillations of the viscera and/or by proprioceptive signals from the limbs impacting the respiratory generator (Iscoe and Polosa, 1976; Bramble and Carrier, 1983; Baudinette et al., 1987; Alexander, 1993; Morin and Viala, 2002; Potts et al., 2005; Giraudin et al., 2012).

Two alleged signatures of hyperpnoea to running exercise have in particular fueled this model. Firstly, breathing augmentation during running is often considered to be inherently proportional to the velocity of repetitive limb movements (Bechbache and Duffin, 1977; Eldridge et al., 1981; DiMarco et al., 1983; Casey et al., 1987). Yet, opposite findings have also been reported (Kay et al., 1975), as well as increased respiratory rates during mental imagery of exercise, i.e. without actual movements (Thornton et al., 2001). Secondly, and this is probably the most controversial aspect, the temporal coordination of breaths to strides (often referred to as the “locomotor-respiratory coupling” or LRC), is commonly highlighted as a conserved feature of hyperpnoea to exercise (Bechbache and Duffin, 1977; Bramble and Carrier, 1983; Alexander, 1993; Corio et al., 1993; Mateika and Duffin, 1995; Lafortuna et al., 1996; Boggs, 2002). However, studies in human participants reported a strong heterogeneity in LRC between individuals, from a constant degree of coupling to no coupling at all (Kay et al., 1975; Bernasconi and Kohl, 1993; Daley et al., 2013; Stickford et al., 2015). The LRC may also be favored by auditory cues (Bernasconi and Kohl, 1993) or by experience (Bramble and Carrier, 1983), arguing for the contribution of multiple factors including external stimuli and training. In quadrupeds, the fewer studies available, essentially on running performant species including rabbits, dogs, cats, and horses (Bramble and Carrier, 1983; DiMarco et al., 1983; Corio et al., 1993; Lafortuna et al., 1996), again revealed various degrees and ratios of locomotor-respiratory coordination, and raised the possibility that faster running gaits (i.e. gallop) may impose a stronger coupling. Therefore, a major source of confound about the coordination of locomotor and respiratory rhythms may owe to the variety of species examined thus far, and to the attendant variability in ambulatory modes and in the contributions of pre-determined (i.e. hardwired) versus secondary (i.e. sensory, volitional, acquired through experience) factors. While the above respiratory-locomotor interactions *can* occur, it is thus far from clear that they are an obligatory feature of hyperpnoea to exercise and are the manifestation of pre-determined circuits that impose on respiratory frequency that of the locomotor movements.

Laboratory mice have become the premier model for investigating complex integrated tasks including adaptive motor control (Benarroch, 2007; Bouvier et al., 2010; Ramanantsoa et al., 2011; Talpalar et al., 2013; Ruffault et al., 2015; Kiehn, 2016; Del Negro et al., 2018) with promising benefits for human health (Amiel et al., 2003; Benarroch et al., 2003; Lavezzi and Matturri, 2008). By being housed and raised in a standardized manner across laboratories, mice should give access to the hardwired manifestation of hyperpnoea to exercise, i.e. with minimal influence of volitional control, variations in external stimuli, or prior experience. Furthermore, mice benefit from a large array of “Omics” tool boxes that allow the manipulation of signals, cell types and neural circuit activity combined to quantitative measurements of behaviors. However, the dynamics of hyperpnoea to exercise had yet to be characterized in this resourceful specie, to pave the way for hypothesis-driven investigations of its cellular and circuit underpinnings. To this aim, we developed a novel method for monitoring inspiratory activity chronically, which we combined with video-tracking of the 4 limbs during unrestrained running on a motorized treadmill operating at different speeds. We reveal that inspiratory frequency augments during trotting by about two-folds to a fixed and stable set point value, irrespective of trotting velocities and of inclination. Yet, respiratory rate was further enhanced during gallop. We also demonstrate the absence at all of temporal coordination of breaths to strides at any speed, intensity or gait. Our work therefore highlights a hardwired mechanism that discretely sets respiratory frequency independently of limb movements but in line with the engaged locomotor program.

## RESULTS

### Chronic electromyographic recordings of diaphragmatic activity in running mice

Examining whether and how the rhythms of limb and breathing movements interact in mice requires to access breathing parameters in freely-moving conditions. As an alternative to measuring ventilation by whole body plethysmography (WBP (DeLorme and Moss, 2002)), which is hardly compatible with displacement movements, we thus extended to mice the use of electromyography (EMG) recordings of the diaphragm (Chang and Harper, 1989; Shafford et al., 2006), the main inspiratory muscle. For this we placed 2 steel wires within the peritoneum to superficially contact the diaphragm, i.e. without passing through the muscle itself (Figure 1A, B; see Materials and Methods for details). Upon full recovery, we first placed EMG-implanted animals in a customized WBP chamber and confirmed that diaphragmatic neurograms correlate with the rising phase (i.e. inspiration) of the plethysmography signal which is an estimate of the respiratory volume (Figure 1C). We used the alternating phases of activity and inactivity of diaphragmatic EMG to define respectively inspiration and expiration and to measure their durations (inspiratory time Ti; expiratory time Te; Figure 1D). The cycle-to-cycle interval (Ti+Te) was used to obtain the instantaneous frequency of each inspiratory burst, leading to respiratory rate. The peak amplitude of integrated EMG signals appeared as an estimate of inspiratory flow obtained as the first derivative of plethysmographic signal (Figure 1C, D). Additionally, these measurements could be repeated over up to 28 days post-surgery without signal degradation (Figure 1E). Importantly, and in contrast to WBP, our EMG recordings provide stable inspiratory activity independently of the animal’s displacement movements and can therefore be performed in freely-behaving mice (Figure 1F). Finally, implanted and connected animals behaved similarly as non-operated mice in an open-field arena (Figure 1G, H) indicating that the implantation did not alter the animal’s ability to move spontaneously nor induced pain.

**Figure 1.**
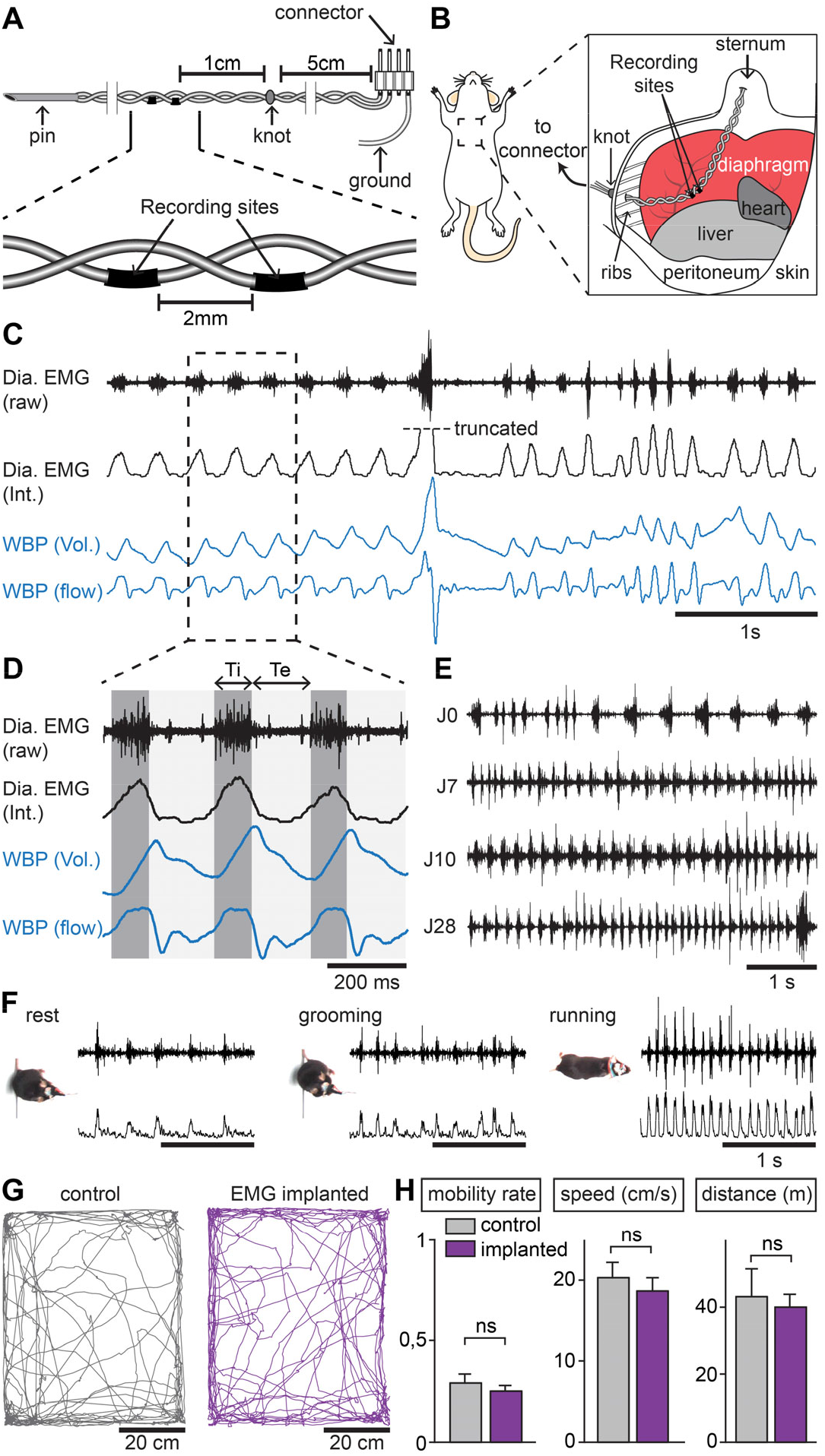
Chronic electromyographic recordings of the diaphragm to monitor inspiration in freely-moving mice. **(A, B)** Schematics of the EMG recording electrodes and of the diaphragm implantation. **(C)** Simultaneous diaphragm EMG recordings (raw and integrated traces in black) and whole-body plethysmography (WBP, volume and flow, in blue) showing that the diaphragm neurogram estimates inspiratory flow. (**D**) Enlarged view of 3 inspiratory bursts highlighting the following respiratory parameters: inspiratory time (Ti) defined as a bout of diaphragm activity, and expiratory time (Te) as the silent period between bouts. **(E)** Diaphragm activity recorded on the day of the surgery (J0) and at 7, 10 and 28 days post-surgery (representative of 4 mice). (**F**) Raw and integrated diaphragm neurograms in an open field test during rest, grooming and spontaneous running showing preserved recording quality in spite of movements. (**G**) Locomotor trajectories of one representative non-implanted mouse (grey) and one representative EMG-implanted mouse (purple) for 10 min in the open-field. (**H**) Bar-graphs showing the mean ± SD mobility rate (left), locomotor speed during mobility (middle) and total distance traveled (right) in control and implanted mice (n=3 in each). ns, not significant; unpaired t-tests.

### Breathing frequency augments during trotting independently of limb velocity

To examine how breathing frequency changes in relation to the velocity of limb movements, we placed EMG-implanted animals on the belt of a motorized treadmill engaged at increasing speeds (25, 40 and 50 cm/s). In this paradigm, mice develop a trotting gait and increase their step frequency linearly to the treadmill speed (Bellardita and Kiehn, 2015; Mayer et al., 2018). We found an immediate increase of respiratory rate at the onset of the animal’s movement, which was surprisingly independent of the treadmill speed (Figure 2A-B). Indeed, respiratory frequency increased by 225% at 25 cm/s, 223 % at 40 cm/s, and by 226% at 50 cm/s, without significant difference between the 3 speeds (Figure 2B). The increase in respiratory frequency was associated with a decrease of both Ti and Te (Figure 2C, D). Additionally, the amplitude of integrated EMG bursts was significantly higher during running than rest, again without significant differences between the 3 speeds tested (Figure 2E). When animals were challenged to a continuous run at 40 cm/s for 10 min, the respiratory frequency was stable from onset throughout the period, excluding an effect of acute stress when the treadmill starts (Figure S1). Secondly, to examine the impact of workload onto breathing changes, mice were submitted to the same running task on a treadmill with a 10% incline ((Gardiner et al., 1982; Gillis and Biewener, 2002), Figure 2F-J). We found that changes in respiratory frequency were similar to the values obtained on the level treadmill and were still independent of running speed (Figure 2G). Furthermore, no differences were found in the changes in Ti, Te nor EMG burst amplitude (Figure 2H-J). Altogether these experiments show that, during trotting in mice, respiratory rate increases to a fixed set point value that is independent of the displacement speed. Furthermore, running on a 10% inclined treadmill did not further enhance breathing rate, suggesting little influence of the exercise grade.

**Figure 2.**
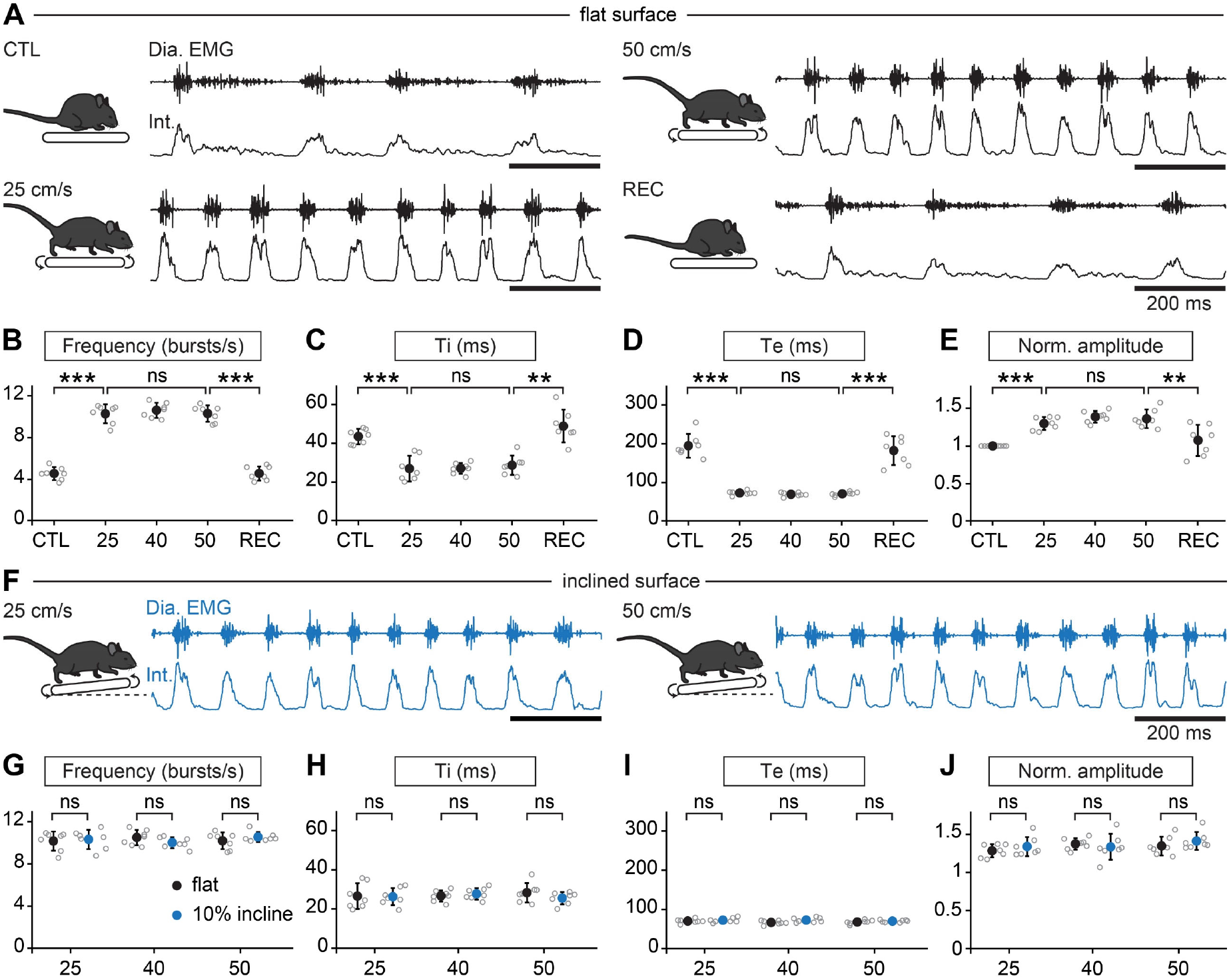
Breathing rate augments during trotting independently of limb velocity or surface inclination. **(A)** Diaphragm activity recordings in control condition (CTL), trotting at 25 and 50 cm/s and during recovery after the run (REC). Raw (Dia. EMG) and integrated (Int.) signals are illustrated in each condition. **(B-E)** Analysis of respiratory parameters in control condition, and during trot at 25, 40 and 50 cm/s, and during recovery: frequency (**B**), inspiratory (Ti, **C**) and expiratory (Te, **D**) times and normalized amplitude (**E**). *** p<0.001; ** p<0.01; ns, not significant; unpaired t-tests. Data are mean ± SD, n=7 mice per condition. **(F)** Raw (Dia. EMG) and integrated (Int.) diaphragmatic activity recordings during trot at 25 and 50 cm/s on a treadmill with 10% incline. **(G-J)** Similar analyses as in (**B-E**) during running on the inclined treadmill. ns, not significant; unpaired t-tests. Data are presented as mean ± SD, n=7 mice per condition. See also Figure S1.

### Breaths are not temporally synchronized to strides during trotting

While our observations above during trotting suggest that the respiratory rhythm can operate without constraints from locomotor movements, a temporal coordination between breaths and strides could occur at specific regimes (Lafortuna et al., 1996). To examine this, each limb was tracked using DeepLabCut ((Mathis et al., 2018), Figure 3A-B, see Materials and Methods) and their coordinates were used to register the time of footfall as well as the stance and the swing phases that define the locomotor cycle ((Bellardita and Kiehn, 2015), Figure 3C). We then expressed the onset of individual inspiratory bursts within the locomotor cycle as a phase value (ΦInsp) from 0 (preceding footfall, FFn-1) to 1 (footfall, FFn, Figure 3C). During trotting on a level treadmill, we found that inspiratory bursts could occur at any moment of the locomotor cycle, regardless of the limb considered as a reference. Indeed, for a given animal, ΦInsp values were evenly distributed across a circular plot diagram (Figure 3D, black marks on the outer circle) resulting in a non-oriented mean phase value (colored dots within the inner circle). Consequently, ΦInsp values collected from all animals were evenly distributed across the entire locomotor cycle (Figure 3D, bar-graphs). Similar observations were made during faster running (50 cm/s), with a 10% incline (Figure 3E) or after trotting for 5 and 10 min (Figure S1D). Altogether, these data demonstrate that, at all regimes of trot, respiratory rate increases without any phasing of breaths to locomotor movements.

**Figure 3.**
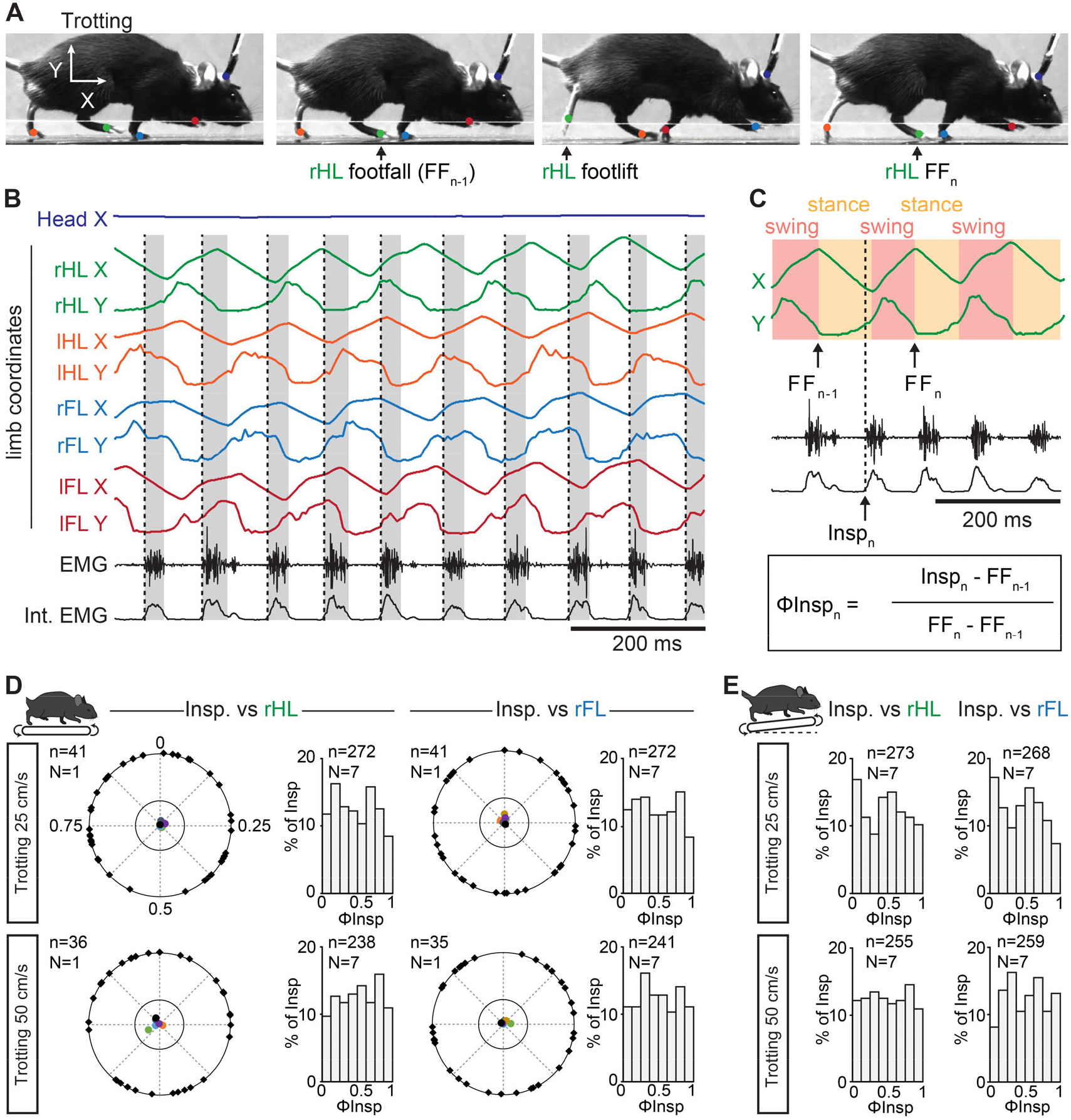
Breaths are not temporally-synchronized to strides during trotting. **(A)** Side views of one representative trotting mouse on a flat treadmill. The 4 limbs and the head were tracked and color-labeled: right hindlimb (rHL, green), left hindlimb (lHL, orange), right forelimb (rFL, blue), left forelimb (lFL, red) and head (dark blue). One complete rHL locomotor cycle is shown, between two consecutive footfalls (FFn-1 and FFn). **(B)** Horizontal (X) and vertical (Y) coordinates of the tracked limbs as well as raw (EMG) and integrated (Int. EMG) diaphragmatic neurograms, during stable trot. Dotted black lines indicate the onsets of inspiratory bursts and shaded rectangles highlight inspiratory times. **(C)** Enlarged view of locomotor cycles of the rHL showing the swing and stance phases. The occurrences of inspiratory bursts (Inspn) within the locomotor cycle are expressed as a phase value (ΦInsp_n_) from 0 (FFn-1) to 1 (FFn). **(D)** Circular plots diagrams showing the phase-relationship between individual inspiratory bursts and the indicated reference limb for one representative animal trotting at 25 or 50 cm/s. Black diamonds on the outer circle indicate the phase of n individual inspirations. The black dot indicates the mean orientation vector for that animal and the colored dots indicate the mean orientation vector of 6 other animals. The positioning of these mean values within the inner circle illustrates the absence of a significantly-oriented phase preference. Bargraphs to the right are distribution histograms of the phases of inspiratory bursts and the same reference limb for all n events from 7 animals. **(E)** Phase distribution histograms between inspiratory bursts and the indicated reference limb for all n events from 7 animals running at 25 or 50 cm/s on the inclined treadmill. Note that inspiratory bursts in (**D**) and (**E**) are evenly distributed across the entire locomotor cycle in each condition and that distribution histograms do not show a phase-preference. See also Figure S1.

### Gallop induces a further augmentation of breathing rate yet without temporal correlation of breaths to strides

While trot allows a wide range of running speeds, quadrupedal mammals including wild-type mice can achieve faster displacement speeds using a galloping gait defined by synchronized movements of the left and right hindlimbs and alternating movements of the left and right forelimbs (Bellardita and Kiehn, 2015; Caggiano et al., 2018; Josset et al., 2018). During gallop in mice, step frequency increases further which entails a greater energetic cost than trot (Heglund and Taylor, 1988; Bellardita and Kiehn, 2015). We hence investigated how breathing rate adjusts to this fast running gait in mice. For this, 7 EMG-implanted animals were placed in a linear runway and a brief air-puff was applied to the back of the animal. In most trials, this evoked a few bouts of gallop (Figure 4A), during which respiratory frequency was found to increase by 314 % from baseline which is significantly higher than the average augmentation measured during trotting (Figure 4B, C). This owed to a further decrease in Ti and Te (Figure 4D, E), and the amplitude of inspiratory bursts was also increased compared to trotting values (Figure 4F). When, following the air puff, animals engaged in trot instead of gallop, their respiratory frequency was significantly lower than at gallop (Figure S2), ruling out that the augmented respiratory rates associated to gallop are influenced by the air stimulus itself. Altogether, these results indicate that the respiratory frequency increase is more pronounced when mice engage in gallop instead of trot, owing to shorter inspiratory and expiratory times.

**Figure 4.**
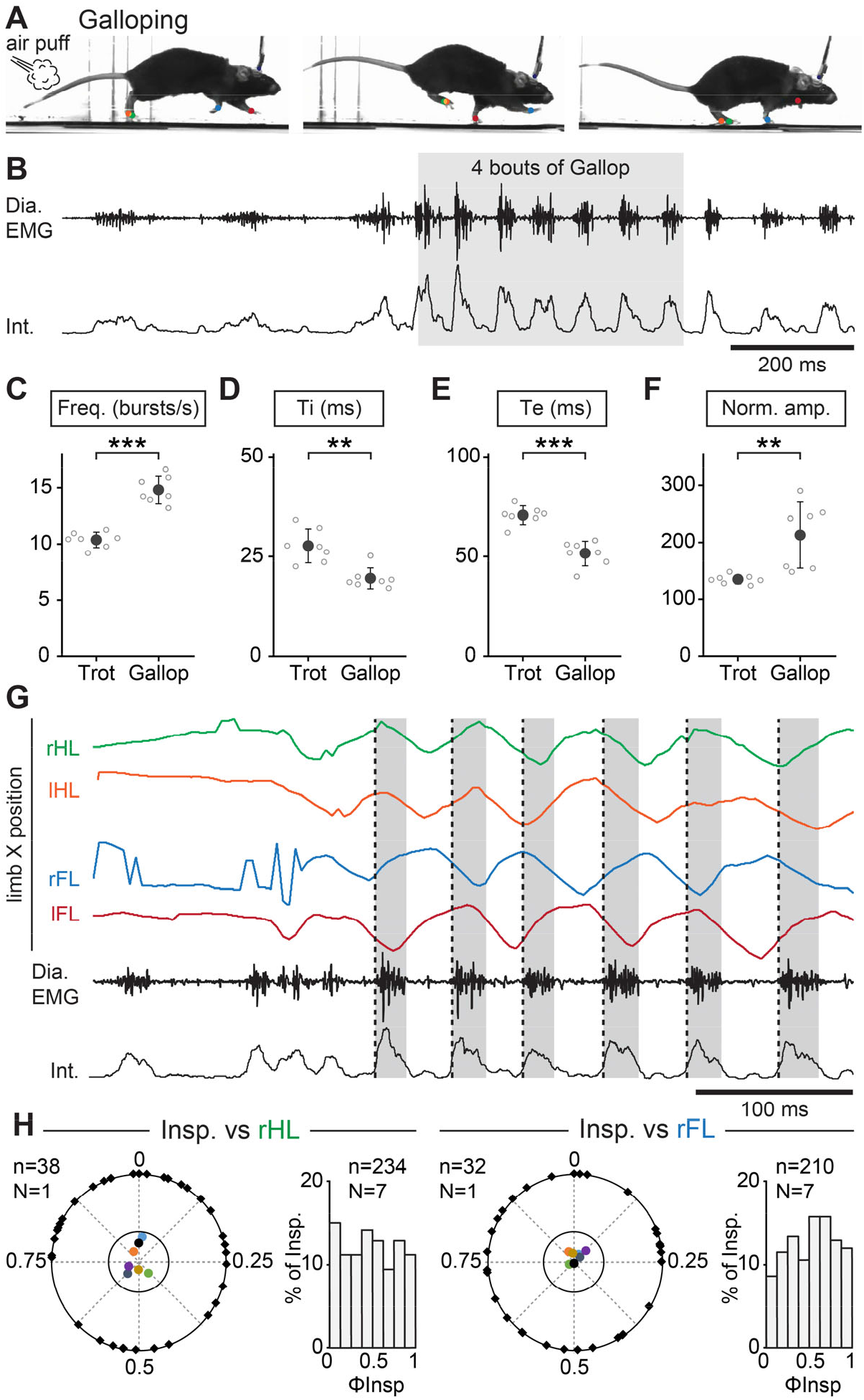
Further augmentation of breathing rate yet without temporal correlation of breaths to strides during gallop. **(A)** Side views of one representative mouse during air puff-induced galloping where right and left hindlimbs (rHL, green; lHL, orange) are synchronized and right and left forelimbs (rFL, blue; lFL, red) are alternating. **(B)** Raw (Dia. EMG) and integrated (Int.) diaphragmatic activity before and during 4 galloping cycles. **(C-F)** Changes in respiratory frequency (**C**), inspiratory (Ti, **D**) and expiratory (Te, **E**) times and normalized amplitude (**F**) between trot and gallop. Note that upon engagement in gallop, breathing rate is further increased as compared to trot. *** p<0.001; ** p<0.01; unpaired t-tests. Data are presented as mean ± SD, n=7 mice for each condition. **(G)** Horizontal (X) coordinates of the 4 limbs as well as raw and integrated diaphragmatic activities before and during 4 bouts of gallop. Dotted black lines indicate the onsets of inspiratory bursts and shaded rectangles highlight inspiratory times. **(H)** Circular plots show the phase-relationship between inspiratory bursts and the indicated reference limb for one representative animal. Black diamonds on the outer circle indicate the phase of n individual inspirations. The black dot indicates the mean orientation vector for that animal and the colored dots indicate the mean orientation vector of 6 other animals. The positioning of these mean values within the inner circle illustrates the absence of a significantly-oriented phase preference. Bar-graphs to the right are distribution histograms of the phases of inspiratory bursts and the same reference limb for all n events from 7 animals.

During gallop, the “hopping-like” movements of the hindlimbs impose a stronger constraint on the visceral mass that may act as a piston mechanism (Baudinette et al., 1987; Alexander, 1993; Bramble and Jenkins, 1993) assisting in producing respiratory airflow. Therefore a constrained occurrence of diaphragmatic movements to a specific phase of the locomotor cycle may be preferred, since possibly most advantageous, at this gait (Baudinette et al., 1987; Lafortuna et al., 1996; Boggs, 2002). We thus examined whether gallop favored a temporal coupling of breaths to strides in mice using the same analytic methods as for trot (Figure 4G). Our data however show that, similar to the trot conditions, breath onsets during gallop were evenly distributed across the locomotor cycle, whether defined using the alternating forelimbs or the left-right synchronized hindlimbs (Figure 4H). Therefore, breaths are not temporally coordinated to strides in mice, even during gallop.

### Prior training does not favor a temporal synchronization of breaths to strides

Since the coordination between breathing and stride rhythms may increase as a function of experience level (Bramble and Carrier, 1983), we next reasoned that some degree of locomotor respiratory synchronization may be acquired through training. Therefore, 3 animals were housed with free access to a running wheel and were trained daily to run on the treadmill for 2 months (Figure S3A), a paradigm known to enhance locomotor skills (Chen et al., 2019; Li and Spitzer, 2020). Animals were then implanted with EMG electrodes and challenged to treadmill running at the 3 speeds on a level or inclined treadmill as well as to the air puff driven gallop as performed above. We observed similar changes in breathing frequency and burst amplitude as in untrained mice (data not shown), and importantly, we found that the onsets of inspiratory bursts were still evenly distributed across the locomotor cycle with no phase preference for any animal and in any condition (Figure S3B-D). Altogether, these experiments and cycle-to-cycle correlation analyses demonstrate that respiratory frequency in mice increases without breaths being temporally-synchronized to locomotor movements, regardless of locomotor speed, grade, gait or prior training.

## DISCUSSION

We provide here the first measurements of breathing during running in the resourceful mouse model, made possible by a unique method for implanting EMG electrodes on the diaphragm combined with limb video tracking. Contrary to WBP, this allows repetitive, long duration and artifact-free recordings of inspiratory activity during locomotor displacements compatible with laboratory testing environments. Compared to implanted nasal probes (Kurnikova et al., 2017) our EMG approach allows a sampling of respiratory activity with much higher temporal resolution. It thus constitutes a timely addition to the toolbox for studying the adaptive control of breathing. Our method was designed to monitor inspiratory activity, the only breathing phase (i) maintained throughout the rest/activity cycle (expiration is passive at rest) and (ii) with a known pacing origin - the preBötzinger complex, the main respiratory rhythm generator (Ausborn et al., 2018; Del Negro et al., 2018). Therefore diaphragm EMGs, if only indirectly reflecting ventilatory flows (Figure 1C), constitute an accurate readout of the temporal organization of sequences of neuronal activity in executive inspiratory circuits. Obviously, the dynamics of lung inflation and deflation also mobilizes abdominal expiratory muscles when switching from passive to active expiration (Abdala et al., 2009), a candidate signature of exercise hyperpnoea. This was only reflected in our data by manifest shortenings of expirations (Te). While other respiratory muscles can be chronically recorded (Romer et al., 2017), the task appears daunting and a source of confounds in the case of abdominal muscles since they may also contribute to posture.

The respiratory changes observed during acute exercise in laboratory mice, housed and raised in a standardized manner across laboratories, make it possible to draw up an outline of the hardwired interactions between respiration and locomotion, i.e. with minimal contribution of intentional control or prior experience. Through that, it will help future hypothesis-driven attempts to identify the long-sought physiological, likely neuronal, substrate for hyperpnoea to exercise (Mateika and Duffin, 1995; Gariepy et al., 2010). On that matter, one major finding of our work is the absence at all, in both naïve or trained mice, of a temporal locking of breaths to strides. Therefore, although a strict temporal locking *can* occur in some subjects (Kay et al., 1975; Bernasconi and Kohl, 1993; Daley et al., 2013; Stickford et al., 2015), species or at specific locomotor regimes (Alexander, 1993; Lafortuna et al., 1996), the complete absence we report in mice and that others reported during swimming in the lamprey (Gravel et al., 2007; Gariepy et al., 2012), makes it unlikely that phasic entrainment of breathing by sensory inputs sensing inertial oscillations of the viscera or limb dynamics (Iscoe and Polosa, 1976; Bramble and Carrier, 1983; Baudinette et al., 1987; Alexander, 1993; Morin and Viala, 2002; Potts et al., 2005; Giraudin et al., 2012) constitutes an hardwired and obligatory component of respiratory adaptation to exercise.

Our data instead favors the quest of a primary mechanism that sets respiratory frequency independently of the locomotor cycle, and even independently of the velocity of limb movements or exercise intensity. Indeed, and this is another major finding, we demonstrate that respiratory rate increased in a step-like manner by about two-folds upon running initiation and was stably maintained thereafter throughout the running period, irrespective of trotting velocities and regardless of difference in height of the route. This is reminiscent of the respiratory changes previously documented during swimming in the lamprey (Gravel et al., 2007) but contrasts, at first glance, with the proportional increase of respiration to limb velocity observed in human participants (Bechbache and Duffin, 1977; DiMarco et al., 1983; Casey et al., 1987) (but see (Kay et al., 1975) for contradictory findings). Yet, these latter studies typically reported ventilatory volumes, which are not accessible with our EMG recordings. Admitted, while our noticing of increased amplitude of diaphragm EMG signals during exercise denotes increased inspiratory muscular efforts and is compatible with increased minute volume ventilation, our method cannot monitor further augmentations of respiratory airflows if supported by additional muscles or inertial movements. Nonetheless, respiratory rates could further increase when animals engaged into gallop as opposed to trot indicating that the respiratory rhythm generator is not operating at maximal frequency regime during trot but rather at a fixed frequency set point. We therefore propose that efficient breathing adaptation to exercise may include, at least in mice, distinct dedicated behavior-dependent solutions. The transition from rest to exercise could engage a default “exercise” breathing at a fixed frequency set point compatible with a range of energetic demands corresponding to ritualistic territorial and explorative tasks and their share of uncertain and imponderable evolutions (change of speed, of slope, etc...). When appropriate, a shift on the fly to gallop or to resting can operate.

The mechanisms by which the frequency set point is achieved, and those allowing transiting from it will need to be investigated. A bulk of evidence indicates that transitions from walk, to trot and gallop are controlled by specific circuits in the brainstem centers for locomotor initiation (Bachmann et al., 2013; Caggiano et al., 2018; Josset et al., 2018) and in the executive centers of the spinal cord (Andersson et al., 2012; Talpalar et al., 2013; Bellardita and Kiehn, 2015; Skarlatou et al., 2020). Therefore, it is likely that a direct modulation of the brainstem respiratory generator by a neuronal drive of central origin, i.e. from the brainstem and/or spinal locomotor centers, may trigger respiratory frequency increase during exercise (Eldridge et al., 1981; Eldridge et al., 1985; Gariepy et al., 2011; Le Gal et al., 2014). Immediate concerns would be to identify candidate trigger neurons, as well as respiratory frequency clamping mechanisms that need to be blind to the grade of trotting locomotor drives and labile upon gait transitioning. This should be achievable using selective interventional manipulations of locomotor and respiratory neuronal types, now both well-defined and accessible with molecular and genetic markers, viral tracers, and dedicated transgenic mice lines (Bouvier et al., 2010; Talpalar et al., 2013; Ruffault et al., 2015; Kiehn, 2016; Del Negro et al., 2018; Skarlatou et al., 2020).

## MATERIALS AND METHODS

### Animals

All experiments were conducted in accordance with EU directive 2010/63/EU and approved by the local ethical committee (authorization 2020-022410231878). Experiments were performed on C57BL6/J mice of either sex, aged 3 months at the time of the EMG implantation and obtained from Janvier Labs (Le Genest-Saint-Isle, France). All animals were group-housed in plastic breeding cages with free access to food and water, in controlled temperature conditions and exposed to a conventional 12-h light/dark cycle. Animals were managed by qualified personnel and efforts were made to avoid suffering and minimize the number of animals.

### Housing and training protocol

Mice were divided in two groups: untrained mice housed in normal environments, and trained mice that had free access to a running-wheel in the cage. Furthermore, trained mice were exercised daily, from the age of 1 month, for 8 consecutive weeks (see Figure S3A) on a custom-made motorized treadmill with adjustable speed range (Scop Pro, France, belt dimensions: 6 cm x 30 cm). Each training session consisted in placing the animals on the stationary treadmill with no incline for 5 min, before they were engaged for a total of 15 min of running at 3 distinct speeds (25, 40 and 50 cm/s, 5 min at each speed). Mice were allowed to rest for 5 min after running before being placed back in their cage. A week after the EMG implantation, both groups were exercised daily for a week using a similar paradigm but for 1 min at each speed. This step was crucial to obtain stable running animals during experimental sessions.

### Diaphragm EMG recordings

#### Fabrication of EMG electrodes

The protocol was inspired by previous work (Pearson et al., 2005). The electrodes were made of Teflon-coated insulated steel wires with an outside diameter of 0.14 mm (A-M systems, ref 793200). For each animal, a 12 cm pair of electrodes was prepared as follows (Figure 1A). Two wires were lightly twisted together and a knot was placed 5 cm from one end. At 1 cm from the knot, the Teflon insulation was stripped over 1 mm from each wire so that the two bared regions were separated by about 2 mm. The ends of the two wires were soldered to a miniature dissecting pin. The free ends of the electrodes, as well as a 5 cm ground wire, were soldered to a micro connector (Antelec). Nail polish was used to insulate the wires at the connector.

#### Surgical implantation of EMG electrodes in the diaphragm

To implant the diaphragm, 3-month old animals were anaesthetized using isoflurane (4 % at 1 L/min for induction and 2 % at 0.2 L/min for maintenance), placed in a stereotaxic frame (Kopf) and hydrated by a subcutaneous injection of saline solution (0.9 %). Their temperature was maintained at 36°C with a feedback-controlled heating pad. The skull was exposed and processed to secure the micro connector using dental cement (Tetric Evofow). The ground wire was inserted under the neck’s skin and the twisted electrodes were tunneled towards the right part of the animal guided by a 10 cm silicon tube of 2 mm inner diameter. The animal was then placed in supine position, the peritoneum was opened horizontally under the sternum, extending laterally to the ribs, and the silicon tube containing the electrodes was pulled through the opening. The sternum was clamped and lifted upwards to expose the diaphragm. A piece of stretched sterile parafilm was placed on the upper part of the liver to avoid friction during movement of the animal and to prevent conjunctive tissue formation at the recording sites. The miniature dissecting pin was pushed through the right floating ribs. The pin was then inserted through the sternum, leaving the bare part of the wires in superficial contact with the diaphragm (Figure 1B). The position of the electrodes was secured on both sides of the floating ribs and sternum using dental cement. The pin was removed by cutting above the secured wires. The peritoneum and abdominal openings were sutured and a head bar was placed on the cemented skull to facilitate animal’s handling when connecting and disconnecting EMG cables during behavioral sessions. Buprenorphine (0.025 mg/kg) was administered subcutaneously for analgesia at the end of the surgery and animals were observed daily following the surgery.

#### Behavioral experiments

Upon a full week of recovery, implanted animals were connected with custom light-weight cables to an AC amplifier (BMA-400, CWE Inc.) and neurograms were filtered (high-pass: 100 Hz, low-pass: 10 KHz), collected at 10kHz using a National Acquisition card (USB 6211) and live-integrated using the LabScribe NI software (iWorxs).

#### Plethysmography recordings

Four unrestrained implanted animals were connected to the AC amplifier and placed inside a hermetic whole-body plethysmography chamber (Ruffault et al., 2015), customized to allow the passage of the EMG cable. Respiratory volume and diaphragm EMG activity were recorded simultaneously over a period of 10 minutes using the LabScribe NI software. Respiratory volume was derived to obtain the respiratory flow (Figure 1C, D).

#### Open field experiments

For analyzing changes in global locomotor activity, control or implanted and connected animals were placed, one at a time, in a custom open field arena (opaque PVC, 60 x 70 cm) without prior habituation and filmed from above during 10 minutes at 20 frames/s using a CMOS camera (Jai GO-5000-C-USB). The open-source software ToxTrac ((Rodriguez et al., 2017), https://sourceforge.net/projects/toxtrac) was used for automated tracking of the animal’s position over time. Following geometric calibration, the following parameters were extracted for each mice (Figure 1H): the mobility rate defined as the percent of time that the animal spent moving above 7,5 cm/s, the average mobility speed defined as the average speed of the animal during mobility (i.e. above 7,5 cm/s) and the total distance travelled per 10-minute recording. Within each category (i.e. control or EMG-implanted) a grand-mean ± SD across mice was then calculated to produce histograms.

#### Running experiments

We used a custom-built treadmill (ScopPro, France) whose speed was remotely adjusted using a USB servo controller (Maestro Polulu). Implanted untrained (n=7) and trained (n=3) animals followed the same running experiments while recording breathing changes. First, we evaluated breathing changes during running at trot, without or with incline. For each speed, the protocol was as follows. Animals were first placed on the stationary treadmill to monitor basal respiration. Animals were then challenged to trot at the lowest speed (25 cm/s) for 1.5 min followed by a 5 min break. The treadmill was then inclined with a 10% slope and animals exercised at the same speed for 1.5 min. This sequence was repeated for the two other speeds (40 and 50 cm/s) with 5 min of rest between trials. At the end of the running experiment, a breathing recovery period was recorded. We also recorded breathing changes throughout a continuous 10 min exercise at 40 cm/s on the motorized treadmill for n=2 untrained and n=2 trained animals. Finally, we evaluated breathing changes during gallop. For this animals were placed in a linear corridor (80 x 10 cm) and brief puffs of air were applied to the back of the animals to trigger fast escape locomotion which comprises a few steps of gallop, as previously described (Talpalar et al., 2013). The test was repeated, with several minutes of rest between trials, until enough gallop bouts (20 to 30 bouts) were acquired.

During running sessions, animals were filmed from the side at 200 fps and 0.5 ms exposure time using a CMOS camera (Jai GO-2400-USB) and images were streamed to a hard disk using the 2^nd^ LOOK software (IO Industries). The start of the EMG recordings was hardware-triggered by the start of the video-recordings, using the frame exposure output of the video camera.

### Automated analysis of limb movements

To track limb movements during running, we used DeepLabCut (version 2.1.5.2, (Mathis et al., 2018), see Figure 3). We manually labelled the positions of the head and the 4 paws from 50 frames of each video. We then used 95% of the labelled frames to train the network using a ResNet-50-based neural network with default parameters for 3 training iterations. We validated with 2 shuffles and found that the test error for trot experiments was: 3.33 pixels and the train error: 2.37 pixels (image size: 1344 x 301). Similarly, we trained the network for gallop condition using a ResNet-50-based neural network with default parameters for 1 training iteration. We validated with 2 shuffles and found that the test error was: 3.43 pixels and the train error: 2.43 pixels (image size: 1936 x 230). These networks were then used to analyze videos from similar experimental settings. X and Y coordinates from the head and the 4 limbs were then extracted and interpolated to 10 kHz to match the EMG recordings. The latter were exported from LabScribe to Clampfit (Molecular Devices), and both sets of signals were merged in a single file, before being processed offline in Clampfit.

### Quantifications and statistics

The instantaneous frequency and amplitude of respiratory bursts were detected using the threshold search in Clampfit from stable trotting moments, i.e. when the animal’s speed was in phase with the treadmill, inferred by the absence of changes in head’s horizontal coordinates. Inspiratory time (Ti) was defined as the duration of the diaphragmatic burst and the expiratory time (Te) as the silent period in between bursts as illustrated in Figure 1D. To ensure homogeneity, measurements were made during 3 stable moments distributed along the recording. Respiratory amplitude change was normalized and expressed as a percent of the control amplitude before running started. Values for respiratory bursts (frequency, Ti, Te and amplitude) were expressed as mean ± SD. Statistical differences between means were analyzed using unpaired Student’s t-tests (GraphPad Prism 7) and changes in respiratory mean values were considered as significant when p<0.05.

## ACKNOWLEDGEMENTS

This work was funded by an ANR grant to JB (ANR-17-CE16-0027) and supported by NeuroPSI, University Paris-Saclay and CNRS. CH holds doctoral fellowships from Région Ile-de-France and Fondation pour la Recherche Médicale. We thank the animal facility housing animals, Aurélie Heuzé for lab management, and Edwin Gatier and Anthony Renard for help with DeepLabCut.

## AUTHOR CONTRIBUTIONS

JB conceived and supervised the study. JB & CH designed experiments. CH & SD performed experiments and analyzed the data. GF contributed to concepts. CH & JB prepared figures. JB wrote the paper with inputs from CH and GF.

## REFERENCES

Abdala, A.P., Rybak, I.A., Smith, J.C., and Paton, J.F. (2009). Abdominal expiratory activity in the rat brainstem-spinal cord in situ: patterns, origins, and implications for respiratory rhythm generation. J Physiol.

Alexander, R.M. (1993). Breathing while trotting. Science 262, 196–197.

Amiel, J., Laudier, B., Attie-Bitach, T., Trang, H., de Pontual, L., Gener, B., Trochet, D., Etchevers, H., Ray, P., Simonneau, M., et al. (2003). Polyalanine expansion and frameshift mutations of the paired-like homeobox gene PHOX2B in congenital central hypoventilation syndrome. Nature genetics 33, 459–461.

Andersson, L.S., Larhammar, M., Memic, F., Wootz, H., Schwochow, D., Rubin, C.J., Patra, K., Arnason, T., Wellbring, L., Hjalm, G., et al. (2012). Mutations in DMRT3 affect locomotion in horses and spinal circuit function in mice. Nature 488, 642–646.

Ausborn, J., Koizumi, H., Barnett, W.H., John, T.T., Zhang, R., Molkov, Y.I., Smith, J.C., and Rybak, I.A. (2018). Organization of the core respiratory network: Insights from optogenetic and modeling studies. PLoS Comput Biol 14, e1006148.

Bachmann, L.C., Matis, A., Lindau, N.T., Felder, P., Gullo, M., and Schwab, M.E. (2013). Deep brain stimulation of the midbrain locomotor region improves paretic hindlimb function after spinal cord injury in rats. Science translational medicine 5, 208ra146.

Baudinette, R.V., Gannon, B.J., Runciman, W.B., Wells, S., and Love, J.B. (1987). Do cardiorespiratory frequencies show entrainment with hopping in the tammar wallaby? The Journal of experimental biology 129, 251–263.

Bechbache, R.R., and Duffin, J. (1977). The entrainment of breathing frequency by exercise rhythm. J Physiol 272, 553–561.

Bellardita, C., and Kiehn, O. (2015). Phenotypic characterization of speed-associated gait changes in mice reveals modular organization of locomotor networks. Curr Biol.

Benarroch, E.E. (2007). Brainstem respiratory control: substrates of respiratory failure of multiple system atrophy. Mov Disord 22, 155–161.

Benarroch, E.E., Schmeichel, A.M., Low, P.A., and Parisi, J.E. (2003). Depletion of ventromedullary NK-1 receptor-immunoreactive neurons in multiple system atrophy. Brain 126, 2183–2190.

Bernasconi, P., and Kohl, J. (1993). Analysis of co-ordination between breathing and exercise rhythms in man. J Physiol 471, 693–706.

Boggs, D.F. (2002). Interactions between locomotion and ventilation in tetrapods. Comparative biochemistry and physiology 133, 269–288.

Bouvier, J., Thoby-Brisson, M., Renier, N., Dubreuil, V., Ericson, J., Champagnat, J., Pierani, A., Chedotal, A., and Fortin, G. (2010). Hindbrain interneurons and axon guidance signaling critical for breathing. Nat Neurosci 13, 1066–1074.

Bramble, D.M., and Carrier, D.R. (1983). Running and breathing in mammals. Science 219, 251–256.

Bramble, D.M., and Jenkins, F.A., Jr. (1993). Mammalian locomotor-respiratory integration: implications for diaphragmatic and pulmonary design. Science 262, 235–240.

Caggiano, V., Leiras, R., Goni-Erro, H., Masini, D., Bellardita, C., Bouvier, J., Caldeira, V., Fisone, G., and Kiehn, O. (2018). Midbrain circuits that set locomotor speed and gait selection. Nature 553, 455–460.

Casey, K., Duffin, J., Kelsey, C.J., and McAvoy, G.V. (1987). The effect of treadmill speed on ventilation at the start of exercise in man. J Physiol 391, 13–24.

Chang, F.C., and Harper, R.M. (1989). A procedure for chronic recording of diaphragmatic electromyographic activity. Brain research bulletin 22, 561–563.

Chen, K., Zheng, Y., Wei, J.A., Ouyang, H., Huang, X., Zhang, F., Lai, C.S.W., Ren, C., So, K.F., and Zhang, L. (2019). Exercise training improves motor skill learning via selective activation of mTOR. Sci Adv 5, eaaw1888.

Corio, M., Palisses, R., and Viala, D. (1993). Origin of the central entrainment of respiration by locomotion facilitated by MK 801 in the decerebrate rabbit. Experimental brain research Experimentelle Hirnforschung 95, 84–90.

Daley, M.A., Bramble, D.M., and Carrier, D.R. (2013). Impact loading and locomotor-respiratory coordination significantly influence breathing dynamics in running humans. PLoS One 8, e70752.

Del Negro, C.A., Funk, G.D., and Feldman, J.L. (2018). Breathing matters. Nature reviews 19, 351–367.

DeLorme, M.P., and Moss, O.R. (2002). Pulmonary function assessment by whole-body plethysmography in restrained versus unrestrained mice. J Pharmacol Toxicol Methods 47, 1–10.

DiMarco, A.F., Romaniuk, J.R., Von Euler, C., and Yamamoto, Y. (1983). Immediate changes in ventilation and respiratory pattern associated with onset and cessation of locomotion in the cat. J Physiol 343, 1–16.

Eldridge, F.L., Millhorn, D.E., Kiley, J.P., and Waldrop, T.G. (1985). Stimulation by central command of locomotion, respiration and circulation during exercise. Respir Physiol 59, 313–337.

Eldridge, F.L., Millhorn, D.E., and Waldrop, T.G. (1981). Exercise hyperpnea and locomotion: parallel activation from the hypothalamus. Science 211, 844–846.

Gardiner, K.R., Gardiner, P.F., and Edgerton, V.R. (1982). Guinea pig soleus and gastrocnemius electromyograms at varying speeds, grades, and loads. J Appl Physiol Respir Environ Exerc Physiol 52, 451–457.

Gariepy, J.F., Missaghi, K., Chevallier, S., Chartre, S., Robert, M., Auclair, F., Lund, J.P., and Dubuc, R. (2011). Specific neural substrate linking respiration to locomotion. Proc Natl Acad Sci U S A.

Gariepy, J.F., Missaghi, K., Chevallier, S., Chartre, S., Robert, M., Auclair, F., Lund, J.P., and Dubuc, R. (2012). Specific neural substrate linking respiration to locomotion. Proc Natl Acad Sci U S A 109, E84–92.

Gariepy, J.F., Missaghi, K., and Dubuc, R. (2010). The interactions between locomotion and respiration. Prog Brain Res 187, 173–188.

Gillis, G.B., and Biewener, A.A. (2002). Effects of surface grade on proximal hindlimb muscle strain and activation during rat locomotion. J Appl Physiol (1985) 93, 1731–1743.

Giraudin, A., Le Bon-Jego, M., Cabirol, M.J., Simmers, J., and Morin, D. (2012). Spinal and pontine relay pathways mediating respiratory rhythm entrainment by limb proprioceptive inputs in the neonatal rat. J Neurosci 32, 11841–11853.

Gravel, J., Brocard, F., Gariépy, J.F., Lund, J.P., and Dubuc, R. (2007). Modulation of respiratory activity by locomotion in lampreys. Neuroscience 144, 1120–1132.

Heglund, N.C., and Taylor, C.R. (1988). Speed, stride frequency and energy cost per stride: how do they change with body size and gait? The Journal of experimental biology 138, 301–318.

Iscoe, S., and Polosa, C. (1976). Synchronization of respiratory frequency by somatic afferent stimulation. J Appl Physiol 40, 138–148.

Josset, N., Roussel, M., Lemieux, M., Lafrance-Zoubga, D., Rastqar, A., and Bretzner, F. (2018). Distinct Contributions of Mesencephalic Locomotor Region Nuclei to Locomotor Control in the Freely Behaving Mouse. Curr Biol 28, 884–901 e883.

Kay, J.D., Petersen, E.S., and Vejby-Christensen, H. (1975). Breathing in man during steady-state exercise on the bicycle at two pedalling frequencies, and during treadmill walking. J Physiol 251, 645–656.

Kiehn, O. (2016). Decoding the organization of spinal circuits that control locomotion. Nature reviews 17, 224–238.

Kurnikova, A., Moore, J.D., Liao, S.M., Deschenes, M., and Kleinfeld, D. (2017). Coordination of Orofacial Motor Actions into Exploratory Behavior by Rat. Curr Biol 27, 688–696.

Lafortuna, C.L., Reinach, E., and Saibene, F. (1996). The effects of locomotor-respiratory coupling on the pattern of breathing in horses. J Physiol 492 (Pt 2), 587–596.

Lavezzi, A.M., and Matturri, L. (2008). Functional neuroanatomy of the human pre-Botzinger complex with particular reference to sudden unexplained perinatal and infant death. Neuropathology 28, 10–16.

Le Gal, J.P., Juvin, L., Cardoit, L., Thoby-Brisson, M., and Morin, D. (2014). Remote control of respiratory neural network by spinal locomotor generators. PLoS One 9, e89670.

Li, H.Q., and Spitzer, N.C. (2020). Exercise enhances motor skill learning by neurotransmitter switching in the adult midbrain. Nat Commun 11, 2195.

Mateika, J.H., and Duffin, J. (1995). A review of the control of breathing during exercise. Eur J Appl Physiol Occup Physiol 71, 1–27.

Mathis, A., Mamidanna, P., Cury, K.M., Abe, T., Murthy, V.N., Mathis, M.W., and Bethge, M. (2018). DeepLabCut: markerless pose estimation of user-defined body parts with deep learning. Nat Neurosci 21, 1281–1289.

Mayer, W.P., Murray, A.J., Brenner-Morton, S., Jessell, T.M., Tourtellotte, W.G., and Akay, T. (2018). Role of muscle spindle feedback in regulating muscle activity strength during walking at different speed in mice. J Neurophysiol.

Morin, D., and Viala, D. (2002). Coordinations of locomotor and respiratory rhythms in vitro are critically dependent on hindlimb sensory inputs. J Neurosci 22, 4756–4765.

Pearson, K.G., Acharya, H., and Fouad, K. (2005). A new electrode configuration for recording electromyographic activity in behaving mice. J Neurosci Methods 148, 36–42.

Potts, J.T., Rybak, I.A., and Paton, J.F. (2005). Respiratory rhythm entrainment by somatic afferent stimulation. J Neurosci 25, 1965–1978.

Ramanantsoa, N., Hirsch, M.R., Thoby-Brisson, M., Dubreuil, V., Bouvier, J., Ruffault, P.L., Matrot, B., Fortin, G., Brunet, J.F., Gallego, J., et al. (2011). Breathing without CO(2) chemosensitivity in conditional Phox2b mutants. J Neurosci 31, 12880–12888.

Rodriguez, A., Zhang, H., Klaminder, J., Brodin, T., and Andersson, M. (2017). ToxId: an efficient algorithm to solve occlusions when tracking multiple animals. Scientific reports 7, 14774.

Romer, S.H., Seedle, K., Turner, S.M., Li, J., Baccei, M.L., and Crone, S.A. (2017). Accessory respiratory muscles enhance ventilation in ALS model mice and are activated by excitatory V2a neurons. Exp Neurol 287, 192–204.

Ruffault, P.L., D’Autreaux, F., Hayes, J.A., Nomaksteinsky, M., Autran, S., Fujiyama, T., Hoshino, M., Hagglund, M., Kiehn, O., Brunet, J.F., et al. (2015). The retrotrapezoid nucleus neurons expressing Atoh1 and Phox2b are essential for the respiratory response to CO(2). Elife 4.

Shafford, H.L., Strittmatter, R.R., and Schadt, J.C. (2006). A novel electrode design for chronic recording of electromyographic activity. J Neurosci Methods 156, 228–230.

Skarlatou, S., Herent, C., Toscano, E., Mendes, C.S., Bouvier, J., and Zampieri, N. (2020). Afadin Signaling at the Spinal Neuroepithelium Regulates Central Canal Formation and Gait Selection. Cell Rep 31, 107741.

Stickford, A.S., Stickford, J.L., Tanner, D.A., Stager, J.M., and Chapman, R.F. (2015). Runners maintain locomotor-respiratory coupling following isocapnic voluntary hyperpnea to task failure. Eur J Appl Physiol 115, 2395–2405.

Talpalar, A.E., Bouvier, J., Borgius, L., Fortin, G., Pierani, A., and Kiehn, O. (2013). Dual-mode operation of neuronal networks involved in left-right alternation. Nature.

Thornton, J.M., Guz, A., Murphy, K., Griffith, A.R., Pedersen, D.L., Kardos, A., Leff, A., Adams, L., Casadei, B., and Paterson, D.J. (2001). Identification of higher brain centres that may encode the cardiorespiratory response to exercise in humans. J Physiol 533, 823–836.

