## Supplemental Figures for "Independent respiratory and locomotor rhythms in running mice"

### SUPPLEMENTAL MATERIAL

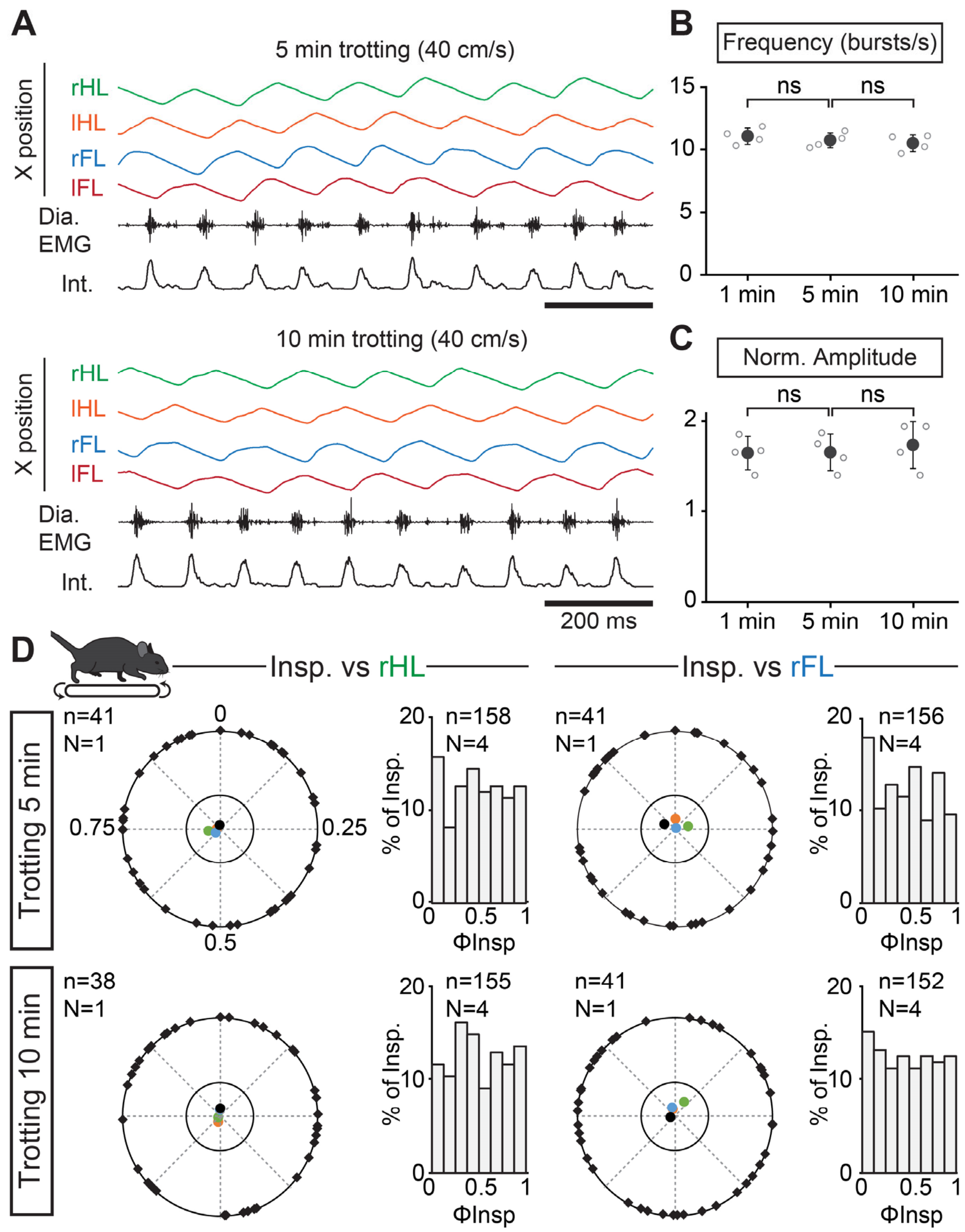


**Supplemental Figure 1. Respiratory changes and correlation with strides during a prolonged treadmill running.**

**(A)** Changes in the horizontal (X) axis of the right hindlimb (rHL, green), left hindlimb (lHL, orange), right forelimb (rFL, blue) and left forelimb (lFL, red) over time, as well as raw (Dia. EMG) and integrated (Int.) diaphragmatic neurograms, after 5 and 10 min of continuous running at 40 cm/s on the level treadmill. **(B-C)** Analyses of respiratory parameters changes after 1, 5 and 10 min of running at 40 cm/s on the level treadmill: frequency (**B**) and normalized amplitude (**C**). ns, not significant; unpaired t-tests. Data are presented as mean ± SD from 4 mice. Note that running for 5 or 10 min did not change the respiratory parameters. **(D)** Circular plots showing the phase-relationship between inspiratory bursts and the indicated reference limb for one representative animal trotting for 5 (top) or 10 min (bottom). Black diamonds on the outer circle indicate the phase of n individual inspirations. The black dot indicates the mean orientation vector for that animal and the colored dots the mean orientation vector of 3 other animals (each in a specific color). The positioning of these mean values within the inner circle illustrates the absence of a significantly-oriented phase preference. To the right of the circular plots are phase distribution histograms between inspiratory bursts and the same reference limb for all n events from 4 animals. Note that all inspiratory bursts are evenly distributed across the entire locomotor cycle in each condition.

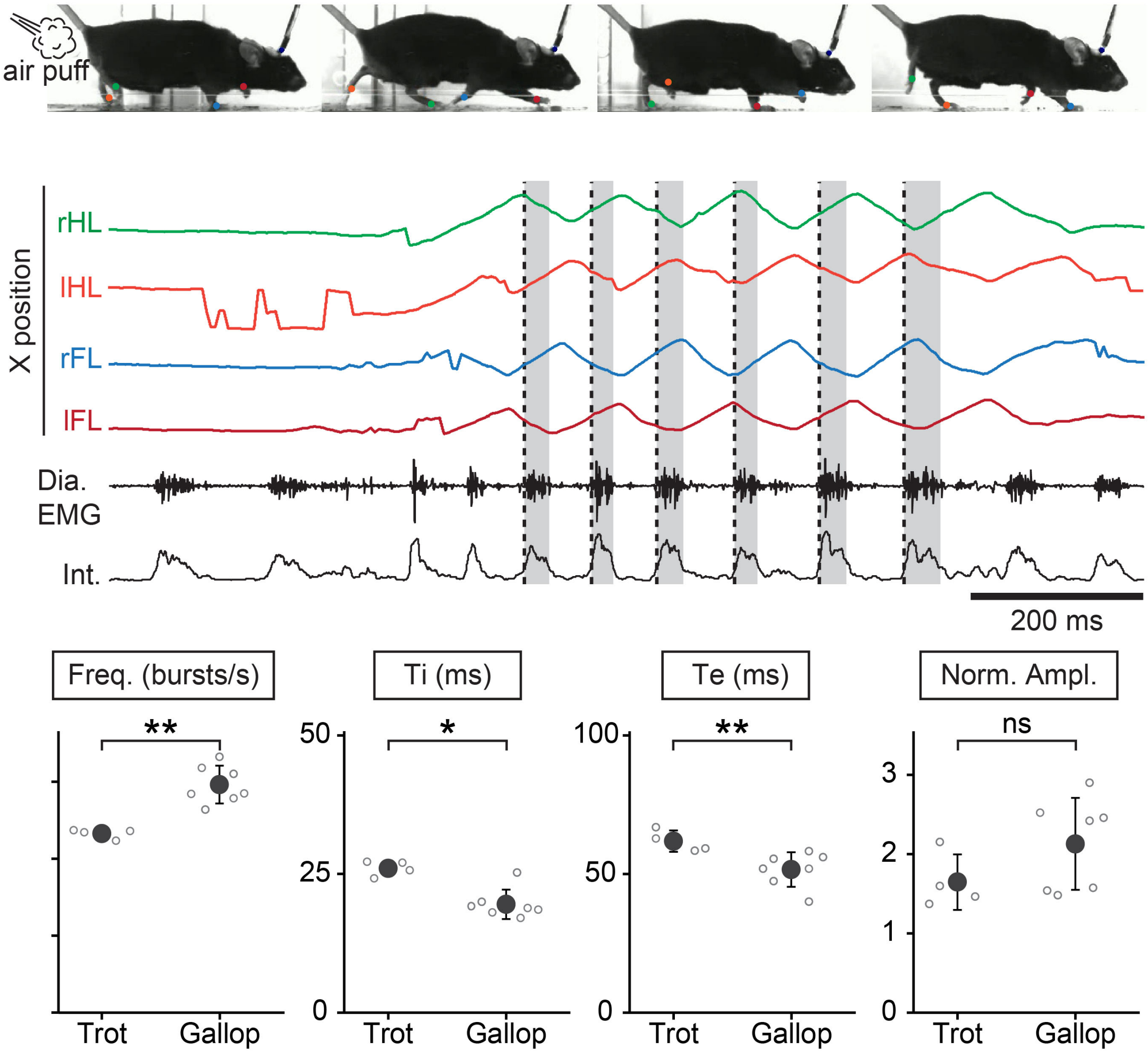


**Supplemental Figure 2. Respiratory changes during air puff-induced trot and gallop.**

**(A)** Side views of one representative mouse during air puff-induced trot where the limbs were tracked using DeepLabCut and color-labeled: right hindlimb (rHL, green), left hindlimb (lHL, orange), right forelimb (rFL, blue) and left forelimb (lFL, red). **(B)** Changes in the horizontal (X) axis of the limbs over time as well as raw (Dia. EMG) and integrated (Int.) diaphragmatic neurograms, before and during 4 cycles of trot induced by air puff. **(C-F)** Analyses of respiratory changes between air puff-induced trot or gallop: frequency (**C**), inspiratory (Ti, **D**) and expiratory (Te, **E**) times and normalized amplitude (**F**). ** $p<0.01$; * $p<0.05$; ns, not significant; unpaired t-tests. Data are presented as mean ± SD, from n=4 mice for trot and n=7 mice for gallop. Note that upon engagement in gallop, breathing is further upregulated as compared to air puff-induced trot.

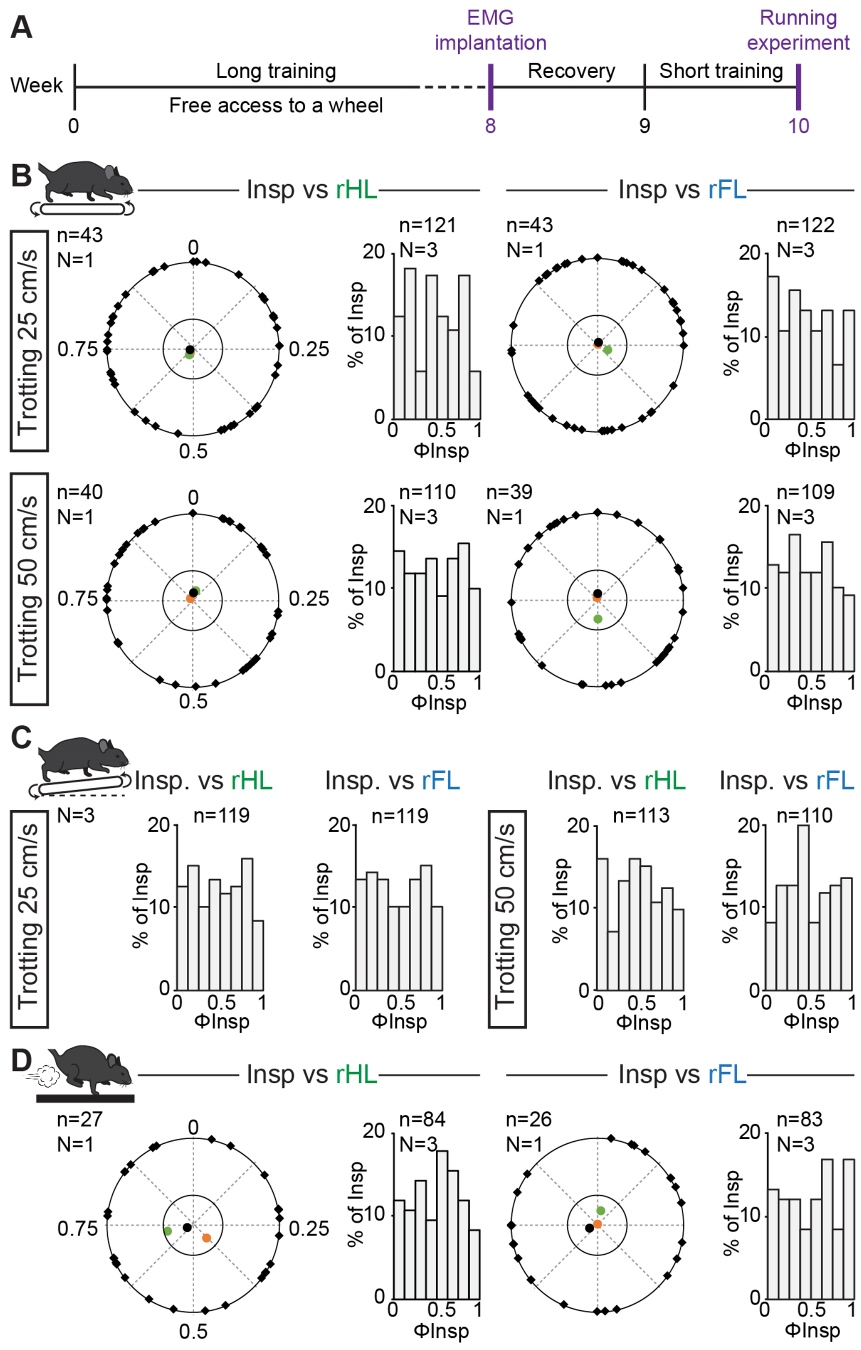


**Supplemental Figure 3. Respiratory changes to exercise in trained mice and phase-relationship analysis between breaths and strides.**

**(A)** Experimental timeline. Mice were trained for 8 weeks before being implanted for EMG recordings of the diaphragm. **(B)** Circular plots showing the phase-relationship between inspiratory bursts and the indicated reference limb for one representative animal trotting at 25 (top) and 50 cm/s (bottom). Black diamonds on the outer circle indicate the phase of n individual inspirations. The black dot indicates the mean orientation vector for that animal and the colored dots the mean orientation vector of 2 other animals. The positioning of these mean values within the inner circle illustrates the absence of a significantly-oriented phase preference. Bar-graphs to the right are distribution histograms of the phases of inspiratory bursts and the same reference limb for all n events from 3 animals. **(C)** Phase distribution histograms between inspiratory bursts and the indicated reference limb for all n events from 3 animals running at 25 cm/s (left) or 50 cm/s (right) on the inclined treadmill. **(D)** Circular plot showing the phase-relationship between inspiratory bursts and the indicated limb for one representative trained animal during gallop. Black diamonds on the outer circle indicate the phase of n individual inspirations. The black circle indicates the mean orientation vector for that animal and the colored circles the mean orientation vector of 2 other animals. To the right are phase distribution histograms between inspiratory bursts and the same reference limb for all n events from 3 animals. Note that all inspiratory bursts are evenly distributed across the entire locomotor cycle in all conditions.
